# FAM19A5 Deficiency Mitigates the Aβ Plaque Burden and Improves Cognition in Mouse Models of Alzheimer’s Disease

**DOI:** 10.1101/2024.05.08.593243

**Authors:** Sumi Park, Anu Shahapal, Sangjin Yoo, Jong-Ik Hwang, Jae Young Seong

**Author notes:** Correspondence and material requests should be addressed to J.Y.S.

## Abstract

FAM19A5, a novel secretory protein highly expressed in the brain, is potentially associated with the progression of Alzheimer’s disease (AD). However, its role in the AD brain remains unclear. Here, we investigated the potential function of FAM19A5 in the context of AD. We generated APP/PS1 mice with partial FAM19A5 deficiency, termed APP/PS1/FAM19A5^+/LacZ^ mice. Compared to control APP/PS1 mice, APP/PS1/FAM19A5^+/LacZ^ mice exhibited significantly lower Aβ plaque density, suggesting that FAM19A5 reduction mitigates Aβ plaque formation. Notably, partial FAM19A5 depletion also prolonged the lifespan of the APP/PS1 mice. To further explore the therapeutic potential of targeting FAM19A5, we developed an anti-FAM19A5 antibody. Administration of this antibody to APP/PS1 mice significantly improved their performance in the novel object recognition test, demonstrating enhanced cognitive function. This effect was reproduced in 5XFAD mice, a model of early-onset AD characterized by rapid Aβ accumulation. Additionally, anti-FAM19A5 antibody treatment in 5XFAD mice led to increased spontaneous alternation behavior in the Y-maze test, indicating improved spatial working memory. These findings suggest that anti-FAM19A5 antibodies may be a promising therapeutic strategy for AD by reducing Aβ plaques and improving cognitive function.

## INTRODUCTION

Alzheimer’s disease (AD) is characterized by the accumulation of amyloid-beta (Aβ) forming plaques in the brain. These plaques are central to the pathogenesis of AD, leading to neuronal damage and cognitive decline[1] Consequently, therapeutic strategies targeting the clearance of Aβ plaques have emerged as a major focus in AD research. These approaches, including anti-amyloid immunotherapies, have shown promise in reducing brain Aβ burden and delaying disease progression[2]. However, clinical trials of anti-amyloid therapies have shown limited effectiveness against cognitive decline, raising concerns about their overall efficacy[3]. Therefore, identifying the ideal target to reverse cognitive decline would be crucial for therapeutic intervention and represents a promising new direction for AD research[4].

FAM19A5, a recently discovered brain-specific protein[5], has emerged as a potential player in both brain development and neurological diseases. Intriguingly, FAM19A5 exhibits a highly localized expression pattern, particularly concentrated in the hippocampus during critical stages of growth[6]. This strategic positioning suggests a role in memory, learning, or other hippocampal functions. Furthermore, recent studies linked FAM19A5 to neurological disorders such as AD, raising the possibility that FAM19A5 level or function contributes to disease pathology[7–10]. It is also known that the progressive loss of synapses in AD is causally related to the decline in memory and cognitive function[11]. Therefore, a recent study suggesting that FAM19A5 may contribute to the process of synapse loss [12] raises the question of whether it is possible to inhibit the pathological progression and cognitive decline of AD using FAM19A5 depletion mice or FAM19A5 antibodies..

In this study, we first generated mice with partial FAM19A5 knockout using the LacZ knock-in technique[6,13]. These mice were then crossbred with a mouse model of AD to study the association between FAM19A5 and amyloid plaque accumulation. We found that partial FAM19A5 deficiency reduced the amyloid plaque burden and extended the lifespan of mice. To develop a potential therapeutic approach, we further generated anti-FAM19A5 antibodies to target FAM19A5 protein in the brain. Administration of these antibodies to AD mice resulted in improved cognitive performance. These results suggest that FAM19A5 could be a promising novel target for therapeutic intervention in AD.

## RESULTS

### Effect of partial FAM19A5 knockout on APP/PS1 transgenic mice

Recent findings suggest that FAM19A5 levels in the brain are potentially correlated with neurodegenerative cascades, including AD[6,9,10]. We hypothesized that reducing FAM19A5 expression would have a therapeutic benefit for AD, leading to improvements in cognitive function. To test this, we generated AD mice with reduced FAM19A5 expression using LacZ knock-in. FAM19A5LacZ mice carry a LacZ reporter gene inserted before exon 4 of the FAM19A5 gene. Consequently, these mice are expected to have lower FAM19A5 levels in the brain than wild-type mice[14].

We then crossed FAM19A5LacZ hemizygous mice with APP/PS1 mice to generate WT, APP/PS1 and APP/PS1/FAM19A5^+/LacZ^ mice. To assess the effects of decreased FAM19A5 levels on Aβ plaque burden, brain tissues from 11-month-old APP/PS1 and APP/PS1/FAM19A5^+/LacZ^ mice were stained with thioflavin-S (Fig. 1A). For image analysis, 6 brain slices from the olfactory bulb to the cerebellum were selected, and the intensity of the signals was measured. Although Aβ plaques were observed in the cortex and hippocampus of both genotypes, APP/PS1/FAM19A5^+/LacZ^ mice exhibited a significantly lower Aβ plaque density than did APP/PS1 mice (Fig. 1B). Partial FAM19A5 depletion also has beneficial effects such as lifespan extension. We tracked the survival rate of APP/PS1 and APP/PS1/FAM19A5^+/LacZ -^ mice for 45 weeks. The survival rate of APP/PS1 mice began to decline at approximately 8 weeks of age, and only approximately 40% of the mice survived to 45 weeks. In contrast, the survival rate of APP/PS1/FAM19A5^+/LacZ^ mice began to decline at approximately 13 weeks of age, and 65% of the mice survived to 45 weeks (Fig. 1C). This result suggested that partial depletion of FAM19A5 may have a protective effect against the decline in survival rates typically observed in the progression of AD.

**Fig. 1.**
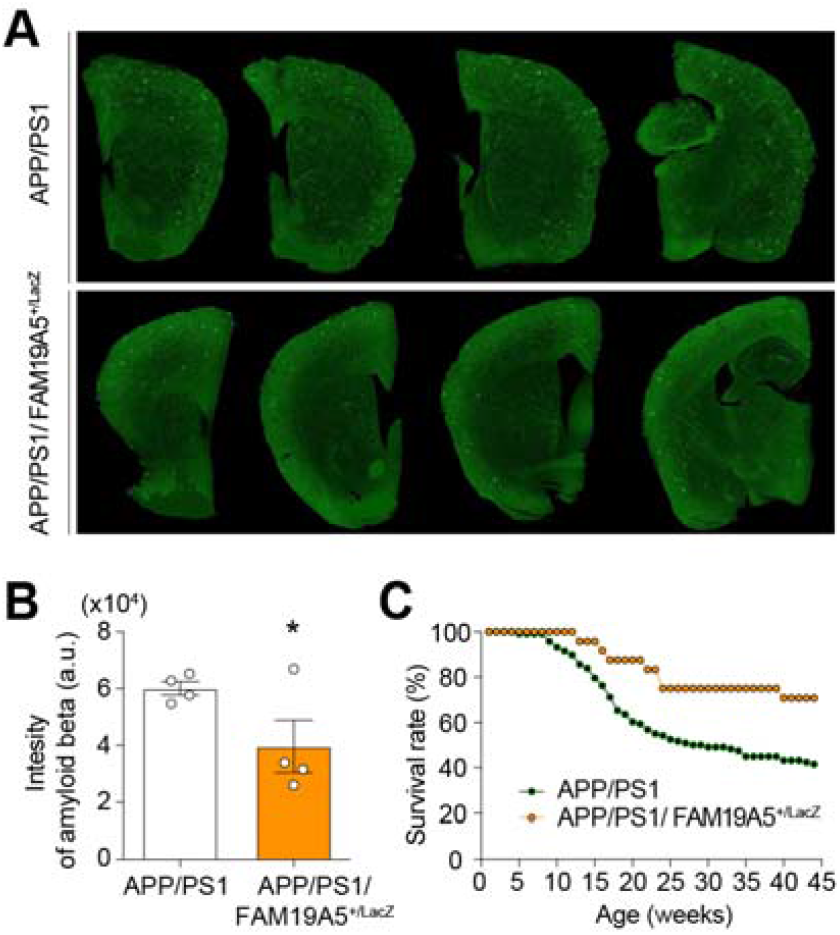
**Aβ** plaque reduction and extended lifespan in APP/PS1/FAM19A5LacZ mice. (A) Thioflavin S staining of brain slices to visualize amyloid plaques (green) and (B) their quantification. Values represent mean ± SEM, n=4 each, Unpaired T test, *p < 0.05. (C) Survival rate of APP/PS1 and APP/PS1/FAM19A5^+/LacZ^ mice, n=118 (APP/PS1), 24

### Effect of the anti-FAM19A5 antibody on cognitive behavior in APP/PS1 mice

The therapeutic effects observed in APP/PS1/FAM19A5^+/LacZ^ mice support the potential of targeting FAM19A5 as a novel therapeutic approach. We developed an anti-FAM19A5 antibody that can be delivered to the brain[12]. We intravenously administered the antibody to APP/PS1 mice and performed a novel object recognition (NOR) test to assess improvements in cognition and memory function. This test consists of 3 phases: habituation, training, and testing. During habituation, the APP/PS1 mice were placed in an open field for 5 min twice a day for 3 consecutive days. Then, the training session was performed in the same open field with two of the same objects for 10 min, and the time spent exploring the object was recorded. All the mice spent similar amounts of time on objects 1 and 2 in the training session. After 6 h, the test session was performed in the same place with one of the objects being replaced by a novel object. The total time spent exploring the novel object and old object was recorded. WT mice spent more time exploring the novel object than the old object. Video tracking during the test session revealed a high preference for the novel object in anti-FAM19A5 antibody-treated mice (Fig. 2A). Human IgG antibody (hIgG) treated APP/PS1 mice, however, did not show a preference for the novel object. Treatment of APP/PS1 mice with an anti-FAM19A5 antibody significantly increased the preference for the novel object (Fig. 2B).

**Fig. 2.**
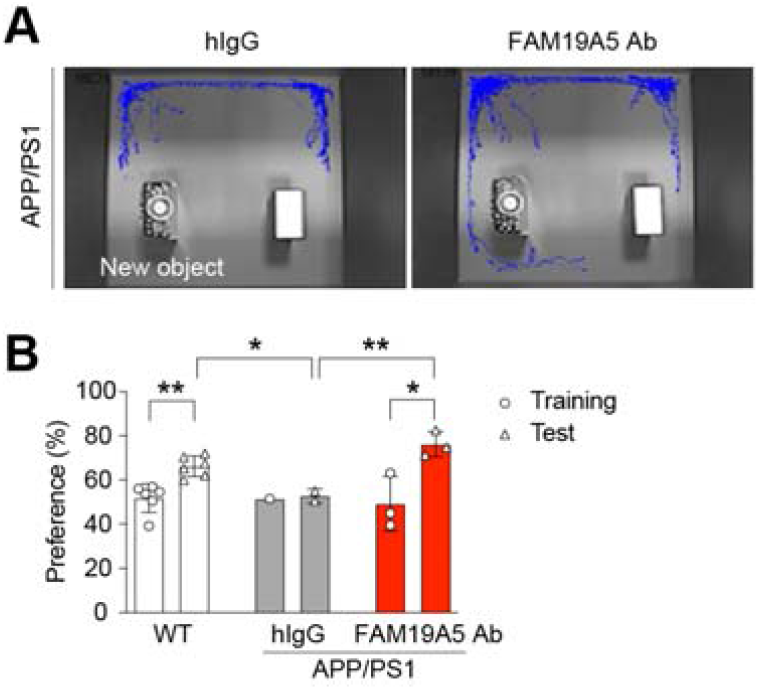
Cognitive improvement in APP/PS1 following anti-FAM19A5 antibody-treatment. (A) Movement trajectories during the test session. (B) Quantification of novel objective recognition after anti-FAM19A5 treatment in APP/PS1 mice. Values represent means ± SEM, n = 1-6, unpaired T test, **p < 0.05, **p

### Effect of the anti-FAM19A5 antibody on cognitive behavior in 5XFAD mice

Given the rapid Aβ accumulation characteristic of the 5XFAD model[15], we employed these mice to assess therapeutic efficacy in a model of early-onset AD. hIgG-treated 5XFAD mice did not exhibit a preference for the novel object in the NOR test, whereas anti-FAM19A5 antibody treatment enhanced exploration of the novel object (Fig. 3A). Across the 3 habituation days, the activity of all mice in the NOR test decreased, as indicated by a main effect of day, suggesting successful habituation. The total distance traveled did not differ among the experimental groups (Fig. 3B).

**Fig. 3.**
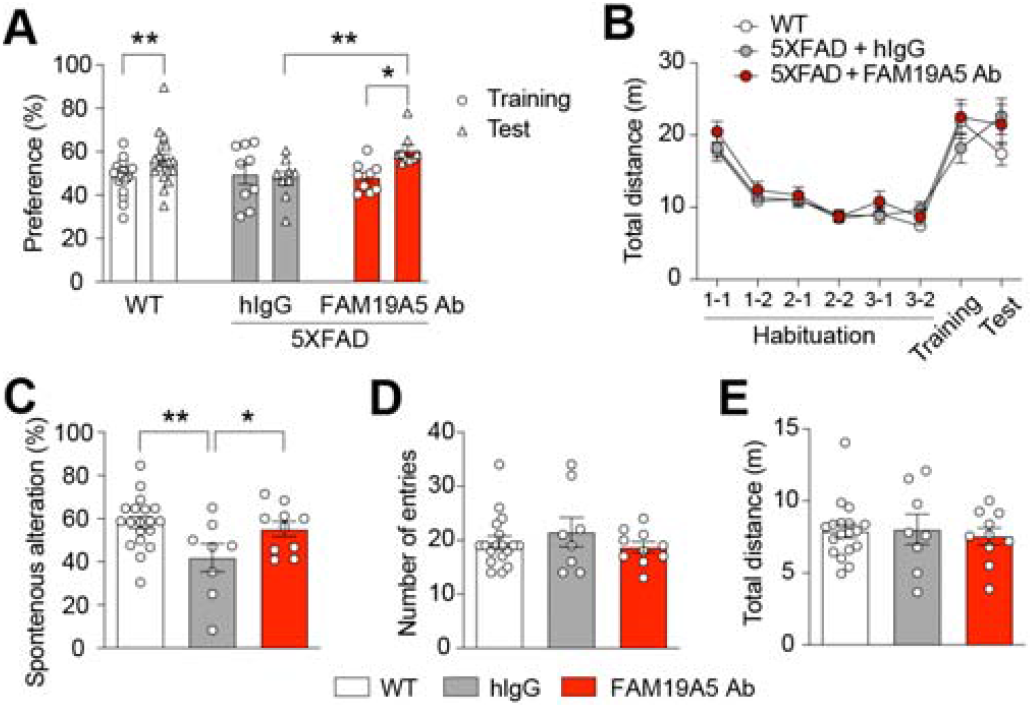
Cognitive improvement in 5XFAD mice following anti-**FAM19A5** antibody-treatment (A) Quantification of novel objective recognition after anti-FAM19A5 treatment in 5XFAD mice. (B) Total traveling distance in meters during the NOR test. n = 8-18, unpaired T test, *p < 0.05, **p < 0.01. (C) Spatial working memory of 5XFAD and WT mice was assessed by spontaneous alternation, (D) total number of arm entries and (E) total distance in the Y maze, respectively. Values represent means ± SEM, n = 8-18, unpaired T test, *p < 0.05, **p < 0.01.

We further assessed spontaneous alternation performance in 5XFAD mice using the Y-maze test. The Y-maze test assesses spatial working memory by measuring the spontaneous alternation rate, which refers to the tendency of the mice to explore a novel arm on each entry. A lower alternation rate is indicative of cognitive impairment (Fig. 3C). Compared with WT mice, 5XFAD mice exhibited significant cognitive impairment. However, compared with 5XFAD control mice, anti-FAM19A5 antibody-treated 5XFAD mice exhibited a significant increase in spontaneous alternations (Fig. 3D). This improvement in exploratory behavior was comparable to that observed in WT mice. Notably, the number of total arm entries and total distance traveled within the maze did not differ between the groups, indicating that motor function and exploratory activity remained normal in 5XFAD mice (Fig. 3E).

## DISCUSSION

In this study, we utilized two well-established AD mouse models, APP/PS1 and 5XFAD, to investigate therapeutic approaches targeting amyloid plaque pathology. Notably, these models exhibit distinct timelines for Aβ plaque formation, with APP/PS1 mice developing plaques at a later stage, mimicking the slower progression of late-onset AD, and 5XFAD mice displaying a more rapid accumulation characteristic of early-onset disease[16]. This distinction highlights the importance of considering disease stage when developing therapies. Our findings in both models suggest that targeting FAM19A5 may be a promising strategy for reducing the Aβ plaque burden. Further investigation is warranted to elucidate the precise molecular mechanisms by which FAM19A5 influences Aβ plaque formation and to determine its therapeutic potential across the AD spectrum.

The present study employed two well-established behavioral tests, the novel object recognition test and the Y-maze, to assess the cognitive benefits of anti-FAM19A5 antibody treatment in APP/PS1 and 5XFAD mice. Notably, treatment with the antibody resulted in identical improvements in cognitive function in both mouse models, as evidenced by their performance in these tasks. This observation is particularly significant because these models represent distinct stages of AD. The identical efficacy across models suggests that targeting FAM19A5 may be a disease-modifying therapeutic strategy, potentially offering benefits not only for symptomatic improvement but also for halting or slowing disease progression. This finding is especially promising considering the current challenges faced in clinical trials targeting amyloid beta plaques. Although these plaques are a hallmark pathology of AD, many such therapies have failed to translate into significant cognitive benefits in human patients[3]. Targeting FAM19A5 could represent a novel and more effective therapeutic approach for AD.

Intriguingly, our study revealed that APP/PS1 mice lacking FAM19A5 exhibited a significantly longer lifespan than wild-type APP/PS1 mice. This remarkable finding suggests that a decrease in FAM19A5 can not only impact cognitive function but also influence overall disease progression in AD models. This observation strengthens the potential of anti-FAM19A5 antibodies as a novel therapeutic strategy. Targeting FAM19A5 may not only achieve symptomatic improvements in cognitive function, as suggested by the electrophysiological and behavioral tests in AD mice[12], but also potentially modulate disease progression and extend lifespan in AD patients. Further investigation is crucial to delineate the precise mechanisms by which FAM19A5 deficiency extends lifespan and to assess the efficacy of anti-FAM19A5 antibodies in promoting longevity in preclinical models with advanced features of human AD pathology. Additionally, studies exploring the safety and efficacy of this approach in aged animals are warranted to inform its potential translation into clinical trials for AD patients.

## MATERIALS AND METHODS

### Animals

Double-transgenic mice (B6. Cg-Tg (APPswe, PSEN1dE9)85Dbo/J) (stock #004462) was obtained from the Jackson Laboratory (Bar Harbor, ME, USA). These mice coexpress the PS1△dE9 mutant form of PS1 and a chimeric mouse-human APP695 strain harboring the Swedish K594N and M595L mutations. Two transgenic genes are controlled by independent mouse prion protein promoter elements, which drive high protein expression in neurons and astrocytes of the CNS.

5XFAD transgenic mice, B6. Cg-Tg (APPSwFlLon PSEN1*M146L*L286V) 6799 Vas/Mmjax was obtained from the Korea Institute of Brain Science. Transgenic mice with mutant human APP (695) with the Swedish (K670N, M671L), Florida (I716V), and London (V717I) familial Alzheimer’s disease (FAD) mutations and human PS1 harboring two FAD mutations, M146L and L286V, were generated. The 5XFAD mice accumulate a high level of amyloid deposition at approximately 1.5 months of age, and plaques spread throughout the hippocampus and cortex by 6 months of age.

The FAM19A5LacZ-KI used in this study was generated by the UC Davis Mouse Biology Program (MBP). These transgenic mice were derived from in-house breeding colonies backcrossed onto a C57BL/6 background. For this study, a new crossbred mouse model, APP/PS1/A5LacZ, was generated by crossbreeding APP/PS1 mice with FAM19A5-LacZ heterozygous mice. The genotypes of all mice were determined by PCR using tail genomic DNA. PCR genotyping was carried out with the following two primers:

All the animals were housed under a 12-12 h light-dark cycle (light phase, 8:00 A.M. to 8:00 P.M.) with standard laboratory diet and water available ad libitum. All animal handling and experiments were conducted with the approval of the Institutional Animal Care and Use Committee (IACUC) of Korea University Medical School.

### Production of anti-FAM19A5 antibody

We generated a chimeric chicken/human monoclonal antibody named 3-2 against FAM19A5 by immunizing chickens with purified recombinant FAM19A5 using a previously described method[13].

### Antibody administration

APP/PS1 and 5XFAD mice were treated with an anti-FAM19A5 antibody or human IgG as a control according to three different treatment protocols, protocols 1, 2, and 3. In protocol 1, 6-to 7-month-old APP/PS1 mice were treated with an anti-FAM19A5 antibody (50 µg/mouse) or hIgG (50 µg/mouse) intravenously every 15 days (twice a month) and sacrificed after the behavioral test. In protocol 2, 3-to 4-month-old 5XFAD mice were treated with an anti-FAM19A5 antibody (100 µg/mouse) or hIgG (100 µg/mouse) intravenously every 15 days (twice a month) and sacrificed. In protocol 3, 3-to 4-month-old 5XFAD mice were first confirmed to have cognitive impairment via the Y-maze test, treated with a humanized anti-FAM19A5 antibody (50 µg/mouse) or hIgG (50 µg/mouse) intravenously every 7 days (4 times a month) and sacrificed after the behavioral test.

### Thioflavin-S staining and plaque quantification

To identify Aβ plaques, brain sections were stained with thioflavin-S solution. The sections were treated with 1% thioflavin-S solution for 10 min and then destained in 70% alcohol for 3 min. Images were acquired with a confocal laser scanning microscope (Leica TCS SP8). Five to eight sections of each transgenic brain were selected for quantification of amyloid plaques using ImageJ software (NIH).

### Novel object recognition (NOR)

The object recognition test was performed as previously described[17]. In the habituation phase, the mice were allowed to explore the open field (30 cm × 30 cm × 30 cm) for 5 minutes twice a day at 6-hour intervals for 3 days. On the 4th day, the training phase, in which the mice were placed in the same open field but with two identical objects placed 5 cm from the wall of the open field, was initiated. Mice were placed in such a way that their heads were opposite to the objects, and then video was recorded for 10 minutes. After 6 hours, the test session was performed. This phase is similar to the training session except that one of the familiar objects is replaced by a new or novel object. During all three phases, both the open field and the objects were cleaned with 70% ethanol and dried before use to minimize error due to olfactory cues. This test was analyzed by using ANY-maze behavioral tracking software.

### Y-maze

The Y-maze is a behavioral test used for assessing memory function to test spontaneous alternation performance. This test utilizes a platform constructed of white, nonreflective plastic and consisting of three arms (arm A, B, and C) oriented at a 120° angle relative to each other with a central triangular area (mid zone). The animals were placed at the end of the first arm and then allowed to freely explore for 5 min. The sequence and total number of arms entered were recorded. The percentage of correct alternations is the number of triads containing entries into all three arms/maximum possible alternations (the total number of arms entered -2) × 100. This test was analyzed by using ANY-maze behavioral tracking software.

### Statistical analysis

The data are expressed as the mean ± SEM. Group means were compared using paired and unpaired Student’s t tests, one-way ANOVA or two-way ANOVA followed by Bonferroni’s multiple comparison test. Statistical analysis was performed with GraphPad Prism 5.0 software. Statistical significance was defined as a p value less than 0.05. The results from behavioral studies, quantitative immunoblot analyses and serum levels were statistically evaluated using Student’s t test, one-way ANOVA or two-way ANOVA followed by Bonferroni’s multiple comparison test. Statistical analysis was performed with GraphPad Prism 5.0 software. Statistical significance was defined as a p value less than 0.05.

## DATA AND MATERIALS AVAILABILITY

The raw data and genetic constructs are available upon request to the authors.

## REFERENCES

[1] Hampel, H., Hardy, J., Blennow, K., Chen, C., Perry, G., Kim, S.H., Villemagne, V.L., Aisen, P., Vendruscolo, M., Iwatsubo, T., Masters, C.L., Cho, M, Lannfelt, L., Cummings, J.L., Vergallo, A. (2021). The amyloid-β pathway in Alzheimer’s disease. Mol Psychiatry 26, 5481–5503.

[2] Golde, T.E., Levey, A.I. (2023) Immunotherapies for Alzheimer’s disease, Science 382 1242–1244.

[3] Zhang, Y., Chen, H., Li, R., Sterling, K., Song, W. (2023) Amyloid β-based therapy for Alzheimer’s disease: challenges, successes and future, Sig Transduct Target Ther 8 1–26.

[4] Long, J.M., and Holtzman, D.M. (2019). Alzheimer disease: an update on pathobiology and treatment strategies. Cell 179, 312–339.

[5] Tang, Y.T., Emtage, P., Funk, W.D., Hu, T., Arterburn, M., Park, E.E.J., Rupp, F., (2004) TAFA: a novel secreted family with conserved cysteine residues and restricted expression in the brain, Genomics 83 727–734.

[6] Shahapal, A., Cho, E.B., Yong, H.J., Jeong, I., Kwak, H., Lee, J.K., Kim, W., Kim, B., Park, H.-C., Lee, W.S., Kim, H., Hwang, J.I., Seong, J.Y. (2019). FAM19A5 expression during embryogenesis and in the adult traumatic brain of FAM19A5-LacZ knock-in mice. Front Neurosci 13, 917.

[7] Díaz de Ståhl, T., Hartmann, C., de Bustos, C., Piotrowski, A., Benetkiewicz, M., Mantripragada, K.K., Tykwinski, T., von Deimling, A., Dumanski, J.P. (2005) Chromosome 22 tiling-path array-CGH analysis identifies germ-line- and tumor-specific aberrations in patients with glioblastoma multiforme, Genes Chromosomes Cancer 44 161–169.

[8] Yilmaz, G., Alexander, J.S., Erkuran Yilmaz, C., Granger, D.N., (2010) Induction of neuro-protective/regenerative genes in stem cells infiltrating post-ischemic brain tissue, Exp Transl Stroke Med 2 11.

[9] Otero-Garcia, M., Mahajani, S.U., Wakhloo, D., Tang, W., Xue, Y.Q., Morabito, S., Pan, J., Oberhauser, J., Madira, A.E., Shakouri, T., Deng, Y., Allison, T., He, Z., Lowry, W.E., Kawaguchi, R., Swarup, V., Cobos, I. (2022) Molecular signatures underlying neurofibrillary tangle susceptibility in Alzheimer’s disease, Neuron 110 2929-2948.e8.

[10] Mathys, H., Davila-Velderrain, J., Peng, Z., Gao, F., Mohammadi, S., Young, J.Z., Menon, M., He, L., Abdurrob, F., Jiang, X., Martorell, A.J., Ransohoff, R.M., Hafler, B.P., Bennett, D.A., Kellis, M., Tsai, L.H. (2019) Single-cell transcriptomic analysis of Alzheimer’s disease, Nature 570 332–337.

[11] Tzioras, M., McGeachan, R.I., Durrant, C.S., Spires-Jones, T.L. (2023) Synaptic degeneration in Alzheimer disease, Nat Rev Neurol 19 19–38.

[12] Kim, H.B., Yoo, S., Kwak, H., Ma, S.X., Kim, R., Lee, M., Ha, N., Pyo, S., Kwon, S.G., Cho, E.H., Lee, S.M., Jang, J., Kim, W., Park, H.C., Baek, M., Park, Y., Park, J.Y., Park, J.W., Hwang, S.W., Hwang, J.I., Seong, J.Y. (2023) Inhibition of FAM19A5 restores synaptic loss and improves cognitive function in mouse models of Alzheimer’s disease, bioRxiv, 2023.11.22.568357.

[13] Kwak, H., Cho, E.H., Cho, E.B., Lee, Y.N., Shahapal, A., Yong, H.J., Reyes-Alcaraz, A., Jeong, Y., Lee, Y., Lee, M., Ha, N., Oh, S., Lee, J.K., Lee, W.S., Kim, W., Hwang, J.I., Seong, J.Y. (2020) Is FAM19A5 an adipokine? Peripheral FAM19A5 in wild-type, FAM19A5 knock-out, and LacZ knock-in mice, 2020.02.19.955351.

[14] Shahapal, A., Park, S., Yoo, S., and Seong, J.Y. (2024). Partial FAM19A5 deficiency in mice leads to disrupted spine maturation, hyperactivity and altered fear response. bioRxiv, 10.1101/2024.04.29.591582

[15] Eimer, W.A., Vassar, R. (2013) Neuron loss in the 5XFAD mouse model of Alzheimer’s disease correlates with intraneuronal Aβ42 accumulation and Caspase-3 activation, Mol Neurodegener 8 2.

[16] Zhong, M.Z., Peng, T., Duarte, M.L., Wang, M., Cai, D. (2024) Updates on mouse models of Alzheimer’s disease, Molecular Neurodegeneration 19 23.

[17] Leger, M., Quiedeville, A., Bouet, V., Haelewyn, B., Boulouard, M., Schumann-Bard, P., Freret, T. (2013) Object recognition test in mice, Nat Protoc 8 2531–2537.

